# Whole-Genome sequencing of Indigenous *Withania somnifera* accession and comparative cytochrome P450 phylogenomics

**DOI:** 10.64898/2026.04.28.721529

**Authors:** Suruchi Gupta, Prashant Misra, Ravail Singh, Manoj Kumar Dhar

## Abstract

Cytochrome P450 monooxygenases (CYP450s) are key oxidative enzymes that diversify plant specialized metabolites and play a central role in the biosynthesis of bioactive withanolides in *Withania somnifera* (L.) Dunal. Despite their importance, genome-wide information on CYP450s in *W. somnifera* has remained elusive. Herein, the first high-quality genome assembly (2.2 Gb, scaffold N50: 47.4 kb) of an Indian *W. somnifera* cultivar was generated using a hybrid Oxford Nanopore-Illumina sequencing strategy. Comparative analysis with the NCBI reference genome revealed moderate SNP and indel variations, reflecting intraspecific genetic diversity. A comprehensive CYP450 catalog was established and analyzed phylogenomically across nine plant genomes, encompassing both withanolide-producing and non-producing Solanaceae and non-Solanaceae species. Unique CYP families (CYP450A, CYP1194, and CYP705A) were detected exclusively in *W. somnifera,* suggesting lineage-specific metabolic innovations, while Solanaceae-restricted (CYP82E/M) and absent (CYP81B, CYP6) lineages highlight taxonomic divergence. Across all analyzed genomes, 36 conserved CYP450 subfamilies, including triterpenoid-associated members, were identified, suggesting a shared oxidative framework adaptable to specialized metabolism. Moreover, potential candidate genes in the triterpenoid pathway, including CYP72A692_1, CYP72A560_4, CYP716A48, CYP724B2, and CYP51G1, were identified through phylogenetic integration with functionally validated triterpenoid-modifying enzymes from other plant species. Gene family evolution analysis further revealed contraction of monoterpenoid-related subfamilies (CYP76A), implying a metabolic shift toward triterpenoid specialization. The comprehensive genome assembly and CYPome of *W. somnifera* offer a valuable resource for functional characterization, evolutionary analysis, and the identification of genes underlying its specialized metabolism. Furthermore, the study advances our understanding of CYP450 diversity and evolution, revealing lineage-specific innovations, conserved subfamilies, and key candidate genes involved in triterpenoid biosynthesis. Together, these findings lay a foundation for future functional studies and pathway engineering aimed at optimizing the metabolic potential of this important medicinal plant.

## Introduction

*Withania somnifera*, a priority medicinal plant recognized by the National Medicinal Plants Board of India, has gained global importance due to its wide range of pharmacological activities. Its therapeutic potential has been primarily attributed to a unique class of triterpenoid steroidal lactones known as withanolides (Afewerky et al., 2021; Alanazi and Elfaki, 2023). These compounds exhibit diverse health benefits and are increasingly in demand in the pharmaceutical and nutraceutical industries, with the market projected to grow at a compound annual growth rate (CAGR) of nearly 10%. However, the natural levels of withanolides in *W. somnifera* remain extremely low (0.001–0.5% dry weight), which poses a major challenge for large-scale utilization. To address this, understanding the genetic and molecular basis of withanolide biosynthesis and elucidation of pathway is of utmost importance. While the early steps of isoprenoid synthesis through the mevalonate (MVA) and methylerythritol phosphate (MEP) pathways are well established, the downstream genes directly involved in withanolide production are still not fully defined. Among them, sterol Δ24-isomerase (24ISO) has been identified as a functionally validated gene that catalyzes a key rearrangement in sterol side chains, regarded as a committed enzyme in withanolide biosynthesis. Although the presence of 24ISO correlates with withanolide production in several Solanaceae species, this link is not universal. For example, some plants such as *Capsicum annum* and *Petunia* carry the 24ISO gene but do not produce withanolides, while species like *Eucalyptus grandis, Senna siamea,* and *Ajuga reptans* synthesize withanolides despite lacking 24ISO gene. This peculiar situation suggests that, although 24ISO is an important gene, its mere presence in the genome is insufficient to classify a plant as capable of producing withanolides. Further the likely involvement of additional enzyme families, particularly cytochrome P450 monooxygenases (CYP450s), which are known to provide pathway specificity by introducing diverse modifications to core metabolite scaffolds, cannot be ruled out (Fig. 1).Moreover, several CYP450 genes in *W. somnifera* and related Solanaceae species are located in close genomic proximity to 24ISO, suggesting their coordinated recruitment in withanolide biosynthesis (Knoch et al., 2018; Priego-Cubero et al., 2025).

**Fig. 1.**
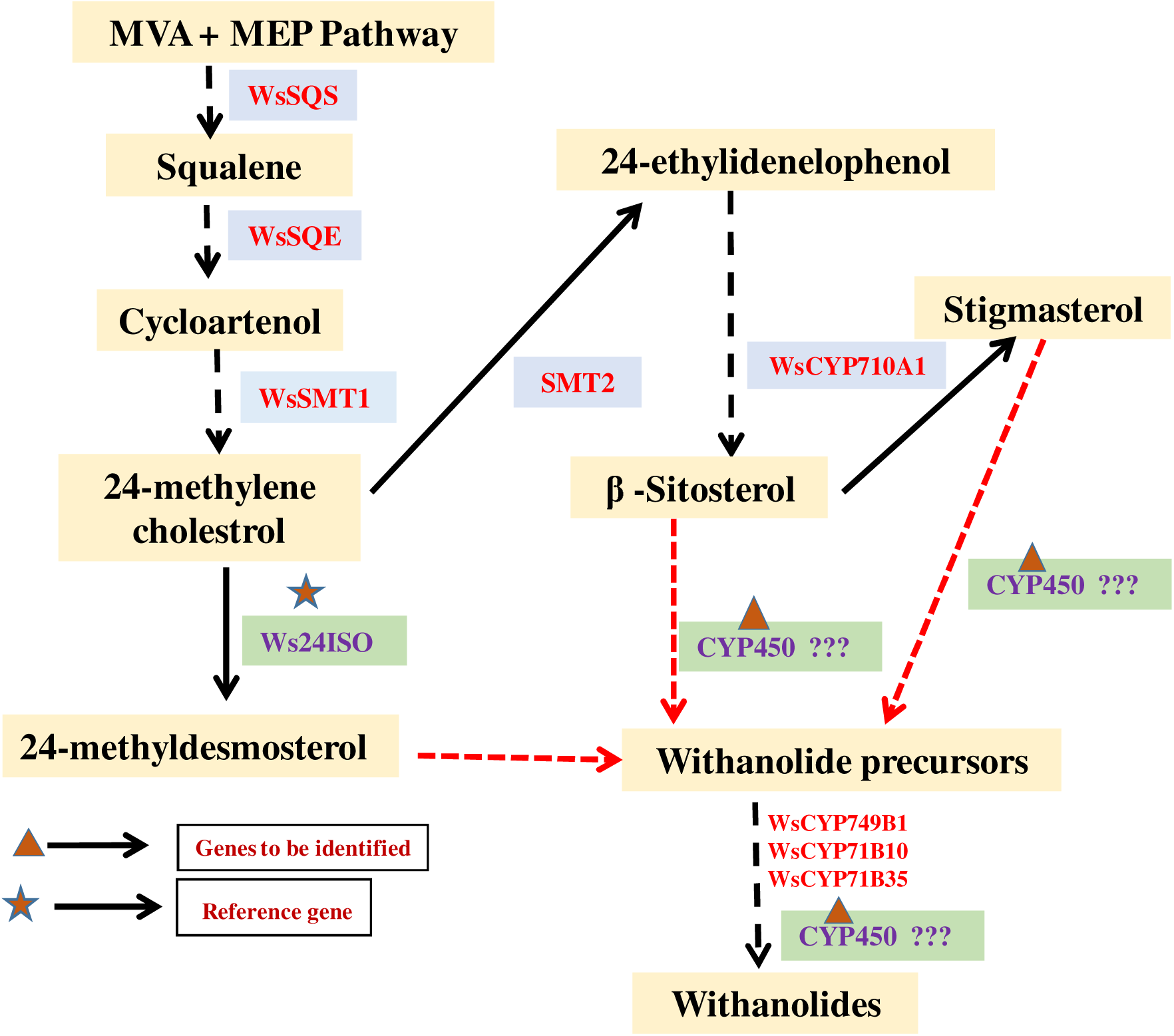
Schematic representation of the proposed role of CYP450 enzymes in the withanolide biosynthetic pathway of *Withania somnifera*

CYP450s represent one of the largest and most versatile enzyme families in plants, catalyzing diverse reactions such as hydroxylation, epoxidation, and oxidation that drive the structural diversification of specialized metabolites. In *W. somnifera*, several CYP450s (including *WsCYP98A,WsCYP76A, WsCYP93Id, WsCYP749B1, WsCYP71B10*, and *WsCYP71B35*) have been functionally characterized and shown to influence withanolide accumulation in a tissue-specific and stress-responsive manner (Rana et al., 2014; Shilpashree et al., 2024; Shilpashree et al., 2022). Earlier our group have also highlighted the functional and regulatory aspects of these enzymes, wherein *WsCYP710A11* was implicated in stigmasterol formation and indirectly in withanolide biosynthesis, exhibiting elicitor-responsive transcriptional modulation (Sharma et al., 2020). Some of these CYP450 genes, including *WsCYP85A, WsCYP90B,* and *WsCYP710A*, are regulated by transcription factors such as WsMYC2, which has been functionally characterized for its positive regulatory role in triterpenoid biosynthesis (Sharma et al., 2019). Despite these advances, the precise P450s involved in scaffold formation and the broader CYPome profile of *W. somnifera* remains to be elusive. Considering the central role of CYP450 diversity in shaping plant secondary metabolism, the present study was conceptualized based on the assumption that differences in CYPome composition may underlie the variable ability of plants to produce withanolides. Therefore, a comparative analysis was carried out across nine plant genomes grouped into three categories: (i) withanolide-producing Solanaceae family members (*Withania somnifera, Physalisgrisea,* and *Lycium barbarum*) (ii) non-withanolide-producing Solanaceae species (*Solanum tuberosum, Solanum lycopersicum,* and *Nicotiana benthamiana*) and (iii) withanolide-producing non-Solanaceae members (*Eucalyptus grandis, Senna siamea,* and *Ajuga reptans)*.

Although CYPome studies have been reported for some Solanaceae species such as *S.melongena* (Lan et al., 2025)*C. annum*(Hao et al., 2022) and *Ipomoea batatas* (Lin et al., 2024), no systematic comparative profiling has been reported across these groups in the context of withanolide biosynthesis. In addition, the present study also entails a comparative analysis between two *W. somnifera* genomes, using the recently published reference assembly retrieved from NCBI (PRJEB64854). This analysis focused on single nucleotide polymorphisms (SNPs) and other sequence differences, providing insights into intraspecific variation that may influence metabolic traits and informing genome-editing strategies.

By and large, this study combines genome-wide CYPome profiling, comparative genomic analysis, and examination of CYP450 family expansion and contraction across withanolide-producing and non-producing taxa. The study reports the first high-quality genome of an Indian *W. somnifera* cultivar that would serves as a valuable platform for functional genomics and genome-editing efforts. The uniquely identified *CYP450* genes associated with triterpenoid metabolism provide strong candidates for further experimental validation, offering opportunities to dissect their roles in withanolide biosynthesis. Additionally, the observed evolutionary patterns of CYP450 families across the examined genomes provide a comparative perspective on how gene family dynamics may contribute to the diversity of specialized metabolites, laying the groundwork for future studies aimed at understanding and harnessing plant metabolic potential.

## 2. Material and Methods

### 2.1 Genome sequencing and Assembly

Seeds of *W.somnifera* were obtained from the CSIR–Indian Institute of Integrative Medicine (CSIR-IIIM) repository, Jammu, India, and sown in the experimental field beds of CSIR-IIIM, Jammu. After four weeks of growth, fresh young leaves were harvested, flash-frozen in liquid nitrogen, and used for genomic DNA extraction with the DNeasy Plant Mini Kit. DNA purity and quality were assessed using a NanoDrop spectrophotometer, and integrity was confirmed on a 0.8% agarose gel. For Illumina sequencing, genomic DNA was enzymatically fragmented, end-repaired, A-tailed, adapter-ligated, and sequenced on the NovaSeq 6000 platform (Illumina, USA) to generate 150 bp paired-end reads, while for Oxford Nanopore Technology (ONT) sequencing, libraries were prepared using the Ligation Sequencing Kit V14 (Oxford Nanopore Technologies, UK) and sequenced on the PromethION platform, with base calling performed using Guppy. Quality assessment of raw Illumina and ONT reads was carried out using FastQC, and filtering and adapter trimming were performed using FastP. Genome size, heterozygosity, and repeat content were estimated by k-mer analysis using Jellyfish and modeled with GenomeScope 2.0. De novo assembly was performed with the GoldFish assembler using ONT long reads, and polishing was achieved with ntEdit through three iterative rounds of error correction with Illumina short reads, followed by scaffolding with the LINKS workflow to improve contiguity. Repetitive sequences were identified with RepeatModeler and masked across the genome using RepeatMasker. Genome completeness was assessed using BUSCO v5.0.0 against the embryophyta_odb10 database to evaluate the presence of universal single-copy orthologs.

### 2.2 Functional Annotation of *W. somnifera* genome

To predict protein-coding genes, a combined evidence-based and *ab initio* strategy was employed. Protein sequences from *Solanum lycopersicum* were used as homology evidence, and TBLASTN searches (E-value ≤ 1e-5) were performed against the assembled genome, followed by cross-validation with DIAMOND BLASTX and filtering with custom scripts to retain high-confidence regions. These regions were aligned to the genome using GenomeThreader, generating preliminary GFF3 annotations, while RNA-seq data were de novo assembled with Trinity and StringTie to provide transcript evidence. Coding sequence and exon features derived from GenomeThreader outputs were converted into AUGUSTUS-compatible hints, with overlapping exons corrected using custom AWK scripts. A new species model (W) was generated in AUGUSTUS, trained with 53,122 high-confidence gene models obtained from cleaned hints, and the final AUGUSTUS pipeline (e-training) was used to produce the predicted set of protein-coding sequences. Functional annotation of predicted proteins was performed by BLASTp alignment against the UniProtKB/Swiss-Prot database, followed by Gene Ontology (GO) and KEGG pathway assignments based on UniProt cross-references.

### 2.3 Comparative analysis of two *W. somnifera* genomes

The *Withania somnifera* genome generated in the present study was compared with the reference genome available at NCBI with accession number PRJEB64854. Repeat content in both assemblies was assessed using repeat-masking approaches, enabling the identification of simple sequence repeats, low-complexity elements, and satellite repeats. The relative abundance of each repeat class was calculated and represented through bar graphs and pie charts. Genome-wide variants were identified by aligning the two assemblies, followed by the detection of single nucleotide polymorphisms (SNPs) and insertions/deletions (indels). High-confidence variants were retained after quality filtering, and SNPs were classified into transitions and transversions to estimate the Ts/Tv ratio. The frequency of singletons and multiallelic sites was also determined to assess levels of genetic diversity. Indels were further analyzed with particular attention to frameshift-inducing mutations that could potentially disrupt gene function.

### 2.4 Identification of CYP450s from the selected genomes

The genome assemblies of the nine selected plant species were retrieved from the NCBI repository, with the exception of *Withania somnifera*, whose genome was sequenced and assembled in-house. Protein-coding sequences were predicted from each genome using AUGUSTUS, and the resulting proteomes were screened for the presence of Cytochrome P450 (CYP450) domains using both the Pfam database (PF00067) and InterProScan. Sequences containing the characteristic P450 domain were extracted as putative CYP450 candidates. To enable systematic classification, a comprehensive CYPome database was constructed by integrating the identified CYP450 sequences with all reported plant CYP450s available in the curated database maintained by Dr. David Nelson. Based on this reference, the CYP450s from the nine genomes were categorized into respective families, subfamilies, and clans following established nomenclature rules (Supplementary file 1 &2). Finally, custom Python scripts were employed to identify common and unique CYP450 families across the selected genomes, thereby facilitating comparative analyses of CYP450 distribution and diversification.

### 2.5 Gene family expansion and contraction patterns by CAFE Analysis

Protein-coding genes were predicted from the nine selected plant genomes using standardized pipelines, and the resulting datasets were subjected to gene family clustering with OrthoFinder to identify orthologous and paralogous relationships across species. The clustered gene families were subsequently analyzed with CAFE (Computational Analysis of Gene Family Evolution) to estimate patterns of gene family expansion and contraction across the phylogeny. The same framework was applied specifically to cytochrome P450 (CYP) genes, which were first identified from each genome through domain-based annotation and compilation into a CYPome dataset. CAFE was then used to assess expansion and contraction events within the CYPome across species, followed by enrichment analysis to determine CYP subfamilies showing significant evolutionary dynamics. Functional information for significantly expanded or contracted CYP families was inferred through annotation against publicly available databases and literature.

### 2.6 Phylogenetic analysis

To investigate the putative functional roles of cytochrome P450 (CYP) genes identified from the *Withania somnifera* genome, phylogenetic analyses were conducted by integrating these sequences with functionally characterized, triterpenoid-related CYPs from other plant species. Protein sequences ranging from 200 to 2000 amino acids were selected for analysis. Multiple sequence alignments were performed using Clustal Omega (v1.2.4), and phylogenetic trees were constructed employing the FastTree (v2.1.11) algorithm. The resulting trees were visualized using iTOL (Interactive Tree of Life, v6). This approach facilitated the inference of the putative functional roles of W. somnifera CYPs based on their clustering patterns with known triterpenoid-related CYPs. Additionally, to examine the evolutionary relationships among CYP genes across diverse plant species, phylogenetic trees were constructed using CYP sequences from nine selected plant genomes. Representative sequences from each genome were chosen to highlight conserved and divergent lineages. Triterpenoid-related CYPs were further categorized into three subfamilies including CYP71, CYP72, and other triterpenoidassociated subfamilies. Separate phylogenetic trees were generated for each subfamily to resolve orthologous and paralogous relationships in detail.

## 3. Results

### 3.1 Genome Assembly and Functional Annotation of *W. somnifera*

Considering the limited genomic resources currently available for *W. somnifera*, the *de novo* genome assembly represents a crucial step toward better understanding the genetic framework and functional biology of this important medicinal plant. The assembly revealed a total of 95,211 contigs, with the largest contig measuring 258,072 bp and an N50 value of 36,926 bp, indicating the average contig length required to cover 50% of the genome. The GC content of the contigs was 37%, consistent with the typical composition observed in plant genomes. Subsequent scaffolding further improved the assembly quality, reducing the number of sequences to 79,168 scaffolds. The largest scaffold measured 58,341,043 bp, while the N50 increased to 47.4 kb, reflecting a higher degree of contiguity. Importantly, the GC content remained stable at 37% in the scaffolded assembly. Overall, the genome size of *W. somnifera*was estimated at 2.2 Gb (Table 1). The completeness of the genome assembly was evaluated using BUSCO which identified 88.2% complete BUSCOs. Of these, 74.1% were present as single copies, while (14.1%) were duplicated. Additionally, (9.0%) genes were fragmented, and 657 (11%) genes were missing from the assembly, likely due to limitations in sequencing or assembly processes. The sequencing efforts were extensive, generating a total of 1,003,522,000 raw reads, equivalent to 150.5 Gb of data. Of these, 953,867,496 reads (95%) were classified as high-quality reads. Oxford Nanopore Technology (ONT) sequencing further contributed 31,715,103 reads, encompassing 64.68 Gb of data, highlighting the robust sequencing depth achieved for this study.

**Table 1:**
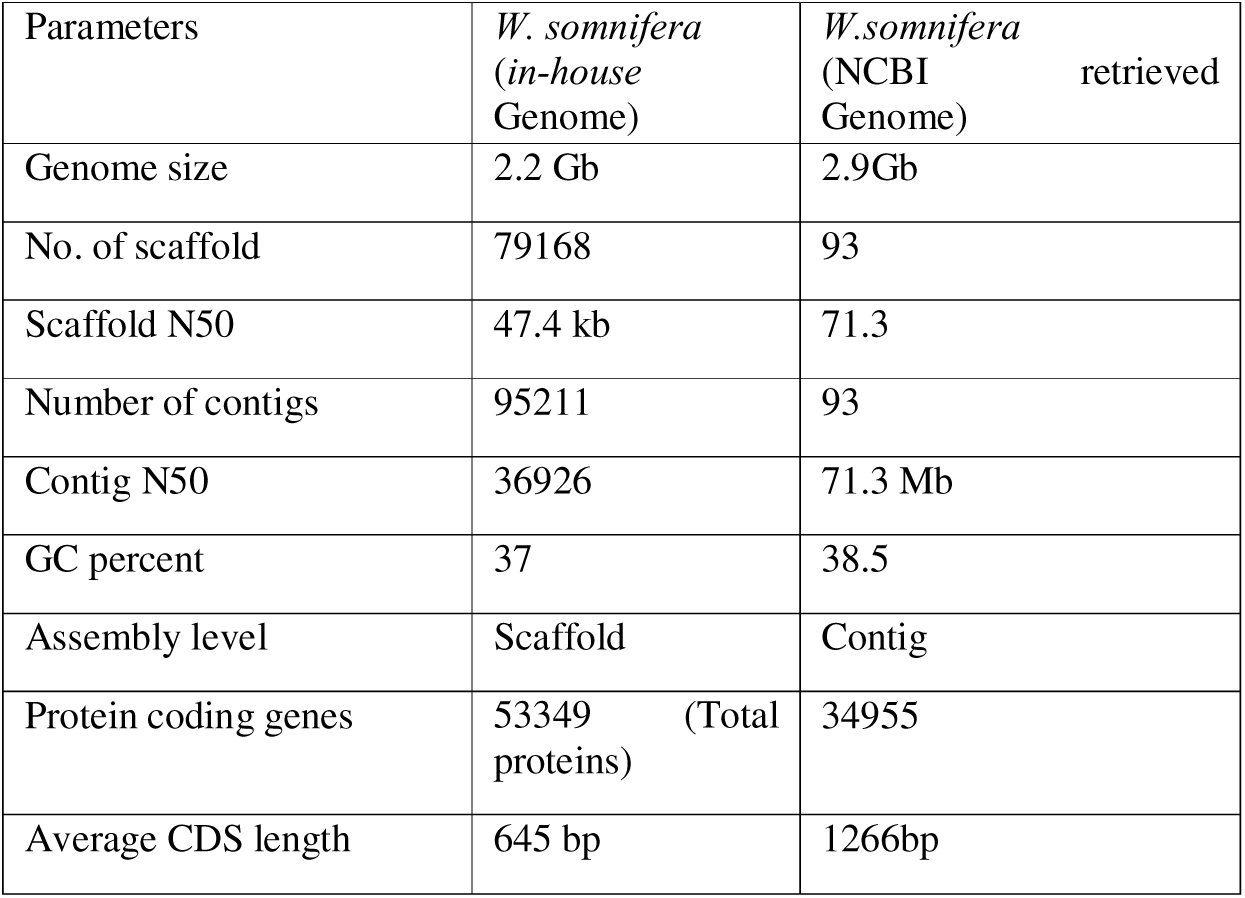
Comparative genome assembly of In-House and NCBI retrieved *W. somnifera* genomes.

Gene prediction in the *W. somnifera* genome using AUGUSTUS identified 31,467 genes, whereas RNA-seq based prediction yielded 47,544 genes. Merging these datasets produced a non-redundant set of 51,198 predicted genes, with an overall completeness of >70%. BLASTx annotation against the UniProt database revealed that 13,596 genes showed significant similarity to known proteins, corresponding to 5,619 unique proteins. Of these, a subset was assigned to COG and KOG orthologous groups, providing insights into their evolutionary relationships and putative functional roles. In addition, 948 proteins were successfully mapped to KEGG pathways, highlighting their involvement in diverse metabolic and signaling networks. Gene Ontology (GO) annotation further classified 14,590 genes, of which 4,672 were associated with biological processes (BP), 5,064 with cellular components (CC), and 4,854 with molecular functions (MF), thereby offering a comprehensive overview of their functional attributes.

The KEGG pathway analysis of proteins identified from the genome of *W. somnifera*revealed that, besides the predominant representation of primary metabolic pathways such as protein modification, amino acid biosynthesis, glycan metabolism, and lipid metabolism, there was notable enrichment in pathways associated with secondary metabolite and isoprenoid biosynthesis (Fig. 2).

**Fig. 2:**
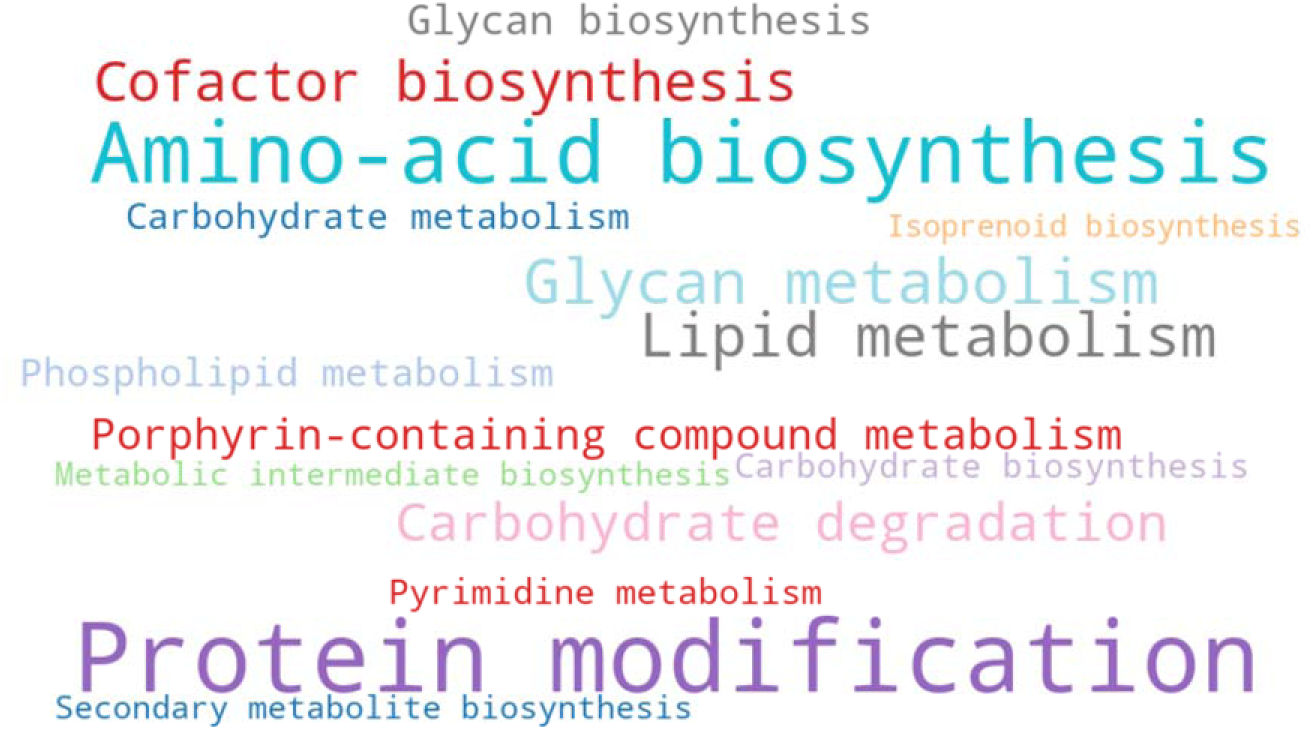
Top 15 KEGG pathways enriched in proteins predicted from the *W. somnifera* genome

GO analysis categorized the predicted proteins into biological process, cellular component, and molecular function. In biological processes, proteins were mainly involved in protein transport, defense response, and protein ubiquitination. Cellular components were enriched for proteins associated with plasma membrane, Golgi apparatus, and chloroplast stroma, indicating important roles in membrane and organelle function. Molecular functions predominantly included ATP binding, metal ion binding, and kinase activities, reflecting active enzymatic and regulatory functions (Fig. 3). Together, these results highlight the functional diversity of the *W. somnifera* proteome, supporting its complex metabolic and regulatory networks essential for growth and secondary metabolite biosynthesis.

**Fig. 3:**
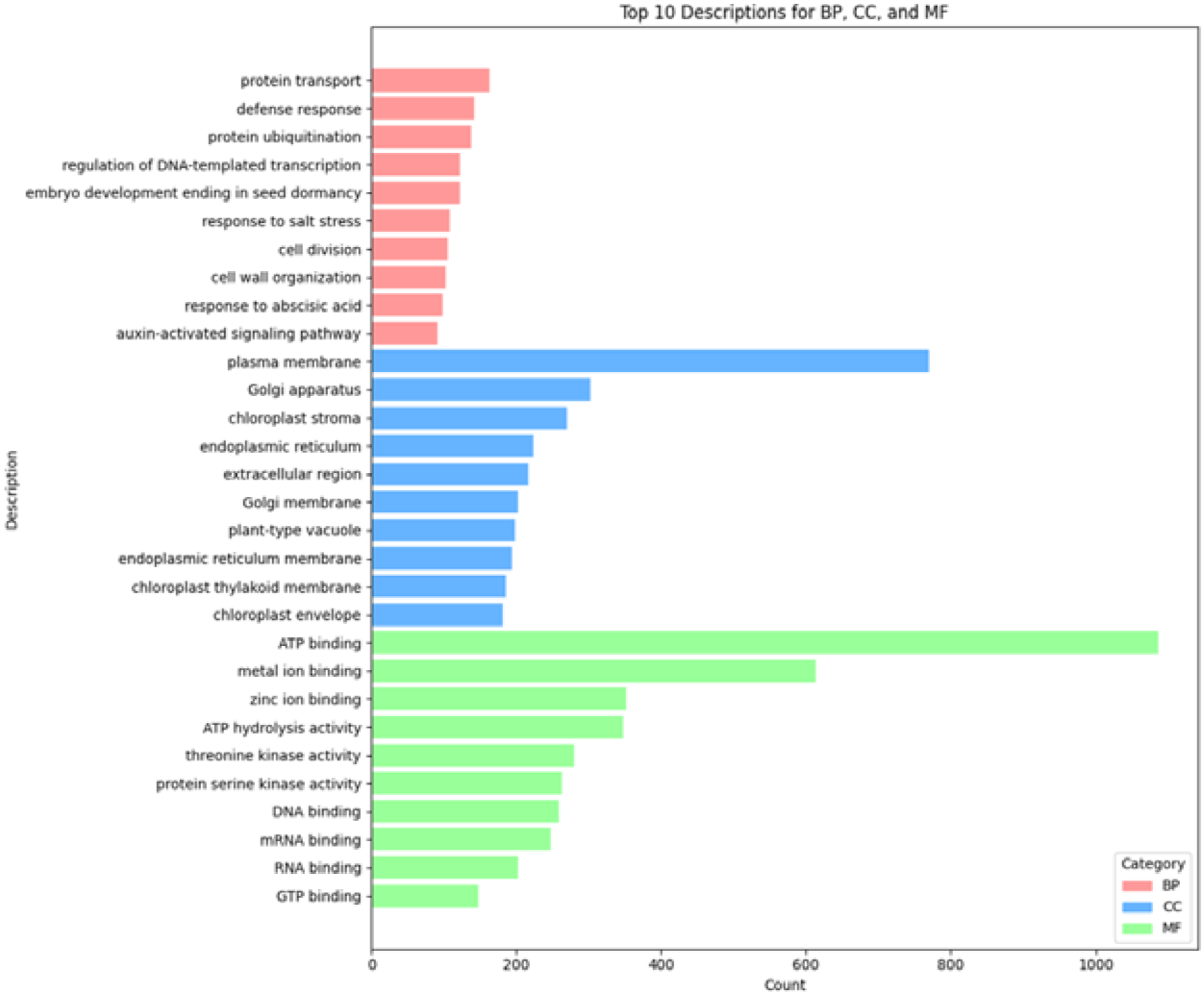
Top 10 Gene Ontology categories of predicted proteins from the *W. somnifera* genome for biological process (BP), cellular component (CC), and molecular functions (MF)

### 3.2 Comparative analysis of two *W. somnifera* genomes

Comparative genome analysis of two *W. somnifera* genomes, an in-house assembly and the reference genome retrieved from NCBI revealed a highly similar repeat composition between the two datasets. Simple sequence repeats were the predominant class, accounting for over 81% of the masked fraction in both genomes, followed by low complexity sequences (∼19%), while satellite repeats were negligible. The overall proportion of masked genome was nearly identical, with only a slight increase in simple repeats observed in the present study. These findings indicate that the current assembly reliably captures the repeat content with high accuracy and consistency relative to the reference genome, thereby supporting the quality of the assembly and annotation (Fig. 4).

**Fig. 4.**
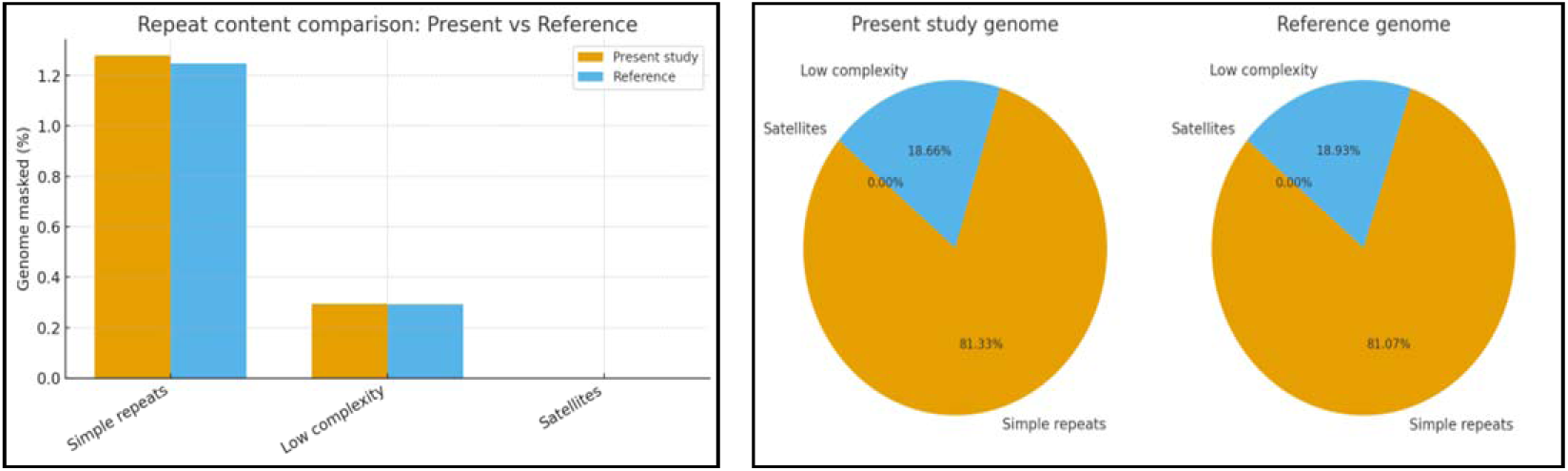
Comparison of repeat content in two *Withania somnifera* genomes. (a) Bar graph showing the proportions of repeat elements. (b) Pie charts depicting the distribution of different repeat classes.

Variant analysis identified 4,491 high-confidence single nucleotide polymorphisms (SNPs) with a transition-to-transversion (Ts/Tv) ratio of approximately 3.2, indicative of reliable variant detection. In addition, 426 insertions and deletions (indels) were observed, constituting about 4% of the total variants, with no frameshift-inducing indels detected, suggesting minimal disruption of gene function. Approximately 14.5% of SNPs were singletons, exhibiting a lower Ts/Tv ratio (1.73), possibly reflecting rare alleles or sequencing artifacts. Furthermore, 37 multiallelic sites, mostly SNPs, highlighted regions with elevated genetic variability (Table 2). Collectively, these results indicate moderate genomic divergence between the two *Withania genomes*, with SNPs representing the primary source of variation and supporting the robustness of the variant dataset for downstream functional and evolutionary analyses

**Table 2:** Genetic Variants Identified in *Withania somnifera* Genome.

| Variant Type | SNPs (%) | ts/tv | Indels (%) | Multiallelic Sites | Multiallelic SNPs |
| --- | --- | --- | --- | --- | --- |
| Singletons | 14.5 | 1.73 | 4 |  |  |
| Multiallelic |  |  |  | 37 | 35 |

### 3.3 Phylogenetics of *W. somnifera* CYP450s with functionally characterized triterpenoid CYPs

Analysis of Cytochrome P450 genes identified from *W. somnifera* genome, together with characterized CYPs involved in triterpenoid pathways from other plant species, revealed subfamily-specific clustering. Further it confirms the accuracy of CYP450 annotation in *W. somnifera* and also highlighted evolutionary associations with characterized plant CYPs, supporting putative roles in sterol and triterpenoid biosynthesis. In addition, several atypical placements were observed, reflecting the divergence events. Overall, the tree is divided into 11 groups, each comprising multiple clades with distinct evolutionary patterns (Fig. 5). Groups 4, 5, 6, 7, 8, and 10 consisted largely of *W. somnifera*-specific sequences, reflecting lineage-specific expansions. Group 4 contained two major clades: one comprising CYP81E and CYP73 members, and another with CYP76 sequences, indicating diversification within these subfamilies. Group 5 was split into two smaller clusters represented by CYP98A and CYP78A families. Group 6 included CYP77 and CYP89 subfamilies, while Groups 7 and 8 were small, containing only a few members (CYP71A, CYP81F, CYP71B20 in Group 7; and CYP736A and CYP83B in Group 8). Group 10 was more complex, comprising five distinct clades dominated by CYP97, CYP96, CYP86, CYP94, and CYP734 families, showing multiphyletic organization.Group 1 was represented primarily by CYP76 members along with two CYP75B genes. *W. somnifera* CYP76B genes clustered closely with previously characterized CYP76s, confirming correct annotation and orthologous relationships. Group 2 contained two major clades: the first comprising CYP93 members, where *W. somnifera* CYP93A3 grouped tightly with CYP93E1 genes from *Glycine max*, *U. uralensis*, *Medicago truncatula*, *Arabidopsis thaliana*, *Vitis vinifera*, and *Quercus suber*, reflecting evolutionary conservation and functional roles in specialized metabolism. The second clade included CYP82 members such as CYP82A4 and multiple CYP82C4 paralogs, which likely originated from gene duplication events and represent functionally diversified variants.Group 3 was divided into four clusters. The first included CYP92A and CYP736A families, suggesting close evolutionary relatedness. The second comprised CYP83B1 paralogs together with the characterized *W. somnifera* CYP71B10, indicating shared motifs and possible functional similarity. The third cluster was dominated by CYP71A2 and CYP71A4 paralogs, while the fourth the largest consisted of CYP71D members, which grouped tightly with the characterized CYP71D from *Nicotiana tabacum*, highlighting conserved functions.Group 9 was composed of four small and one large cluster. The first included CYP51G genes, along with CYP51H10 from *Avena strigosa* and CYP710A1 from the *W. somnifera*. Their close placement reflects conserved roles in sterol biosynthesis, where CYP51 acts as a lanosterol 14α-demethylase and CYP710A1 functions as a sterol C-22 desaturase. The second cluster contained CYP88A members, including *Arabidopsis* CYP88A3, a known triterpenoid enzyme. The third was dominated by CYP707A, while the fourth contained *W. somnifera* CYP724B2 together with CYP708A members from *Q. suber* and *A. thaliana*, suggesting functions in triterpenoid pathways. The largest cluster was dominated by CYP716 members (CYP716A/B/C), which grouped with characterized triterpene-oxidizing enzymes from other plants, reinforcing their probable roles in withanolide biosynthesis.Group 11 was divided into three multiphyletic clades. The first contained CYP7114 and CYP734 members. The second comprised CYP749B members from *W. somnifera*, together with a transcriptome-characterized CYP749B and CYP721A1711. CYP749B family members are implicated in oxidation of fatty acid derivatives and triterpenoid precursors, while CYP721A1711 is associated with phytosterol biosynthesis. Their co-clustering suggests gene duplication followed by neofunctionalization, leading to diversification of functions. The third clade was dominated by CYP72A members. Multiple *W. somnifera* CYP72A genes clustered tightly with characterized CYP72As such as CYP72A154 and CYP72A63, which play critical roles in triterpenoid saponin, glycyrrhizin, and avenacin biosynthesis. This indicates that *W. somnifera* CYP72As are strong candidates for involvement in withanolide and related triterpenoid biosynthesis. The phylogenetic analysis of cytochrome P450 genes from the *W. somnifera* genome, in comparison with well-characterized triterpenoid-related CYPs from other plant species, revealed a set of strong candidate genes potentially involved in withanolide and broader triterpenoid biosynthesis. Particularly, *W. somnifera* members clustered tightly with functionally validated CYP450s from diverse plants, reinforcing their probable roles in scaffold oxidation, precursor modification, and downstream tailoring steps of the triterpenoid pathway. Among them CYP72A554, CYP724B2, CYP716A48, and CYP721A71 showed strong phylogenetic associations with triterpenoid-related CYP clades, suggesting their likely roles in withanolide biosynthesis and making them priority targets for functional validation.

**Fig. 5:**
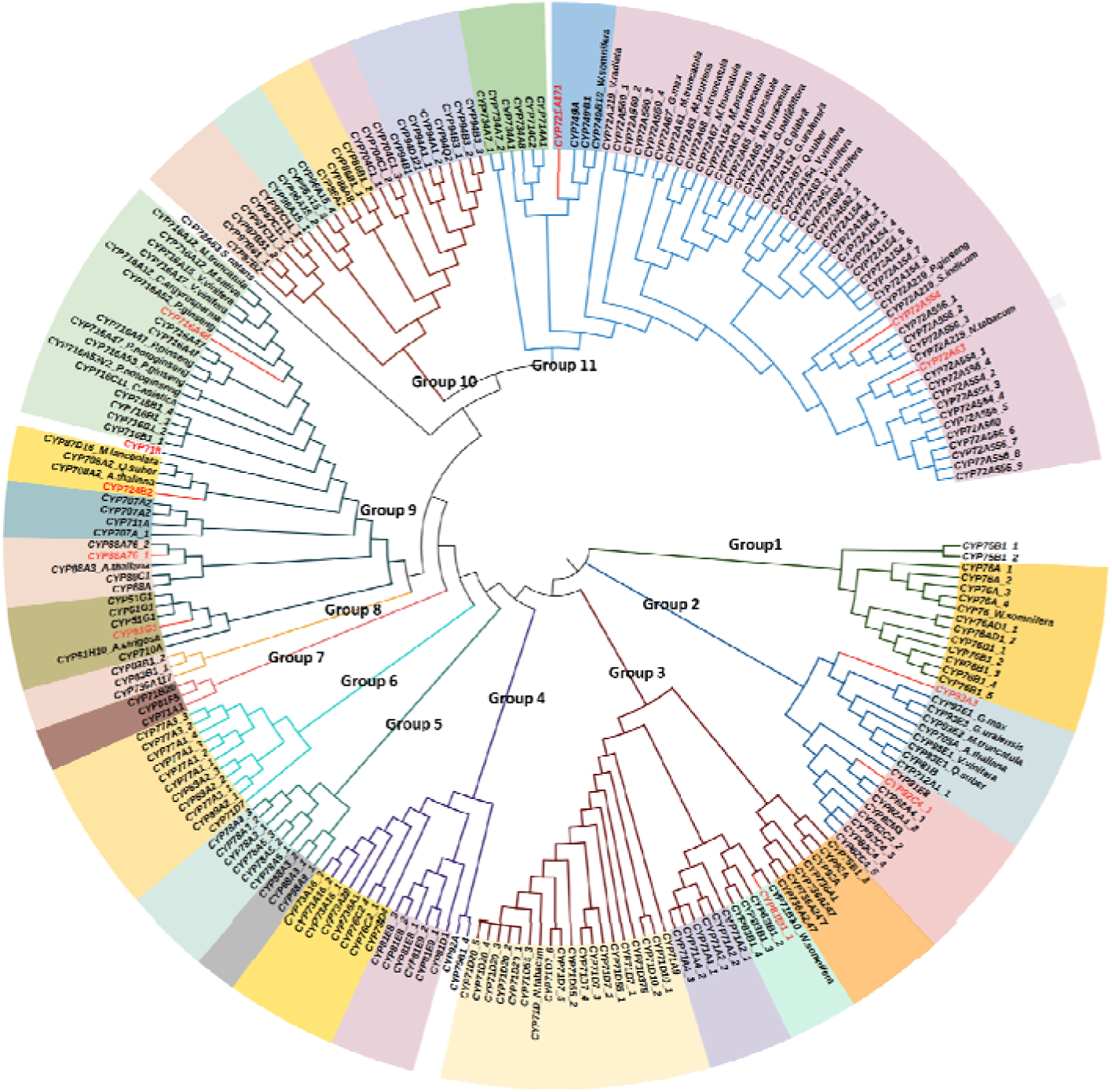
Phylogenetic tree of *W. somnifera* CYP450s showing evolutionary divergence and candidate genes for specialized metabolism. Clades are color-coded to represent groups of CYP450s belonging to similar families/subfamilies

### 3.4 Comparative Genome analysis of selected plant genomes for CYP450 identification

Comparative analysis of CYP450 families across nine plant genomes revealed that CYP450A, CYP1194, CYP80F, CYP82N, CYP84, CYP90, and CYP705A were unique to *W. somnifera*. CYP712A was found only in *W. somnifera* and *Eucalyptus grandis*. CYP6 and CYP81B families were absent from Solanaceae species lacking withanolides, while CYP82E and CYP82M occurred exclusively in Solanaceae genomes. In addition to these unique and lineage-specific families, several CYP450 subfamilies were commonly shared among all nine genomes (Fig. 6). Among these, a total of 36 CYP450 subfamilies were identified, representing conserved clans integral to plant metabolism (Table 3). Representative sequences from each genome and subfamily were used to construct a phylogenetic tree, which resolved into clades consistent with established plant CYP450 phylogeny (Supplementary Fig. 1). The majority of CYPs (243 of 250) formed orthologous subclades across species, reflecting conserved biochemical roles. However, seven CYPs showed deviations from expected orthologous placements, including *W. somnifera* CYP75B and CYP734A, *S. lycopersicum* CYP75B, *N. benthamiana*CYP716A, *S. tuberosum* CYP71A, and *S. siamea* CYP71A, indicating lineage-specific divergence or paralogous relationships. Families such as CYP82C, CYP84A, CYP94A, CYP734A, CYP716A, and CYP78A exhibited strong interspecies conservation, maintaining expected orthology across the analyzed genomes.Analysis of the nine genomes further revealed that CYP71A, CYP72A, CYP93A, CYP76A, CYP94B, CYP81E, CYP88A, and CYP79D are among the commonly shared CYP450s. These genes are associated with triterpenoid pathways and are well-known for catalyzing oxidation, hydroxylation, and other modifications of triterpenoid scaffolds and their derivatives (Supplementary Table 1)

**Fig. 6:**
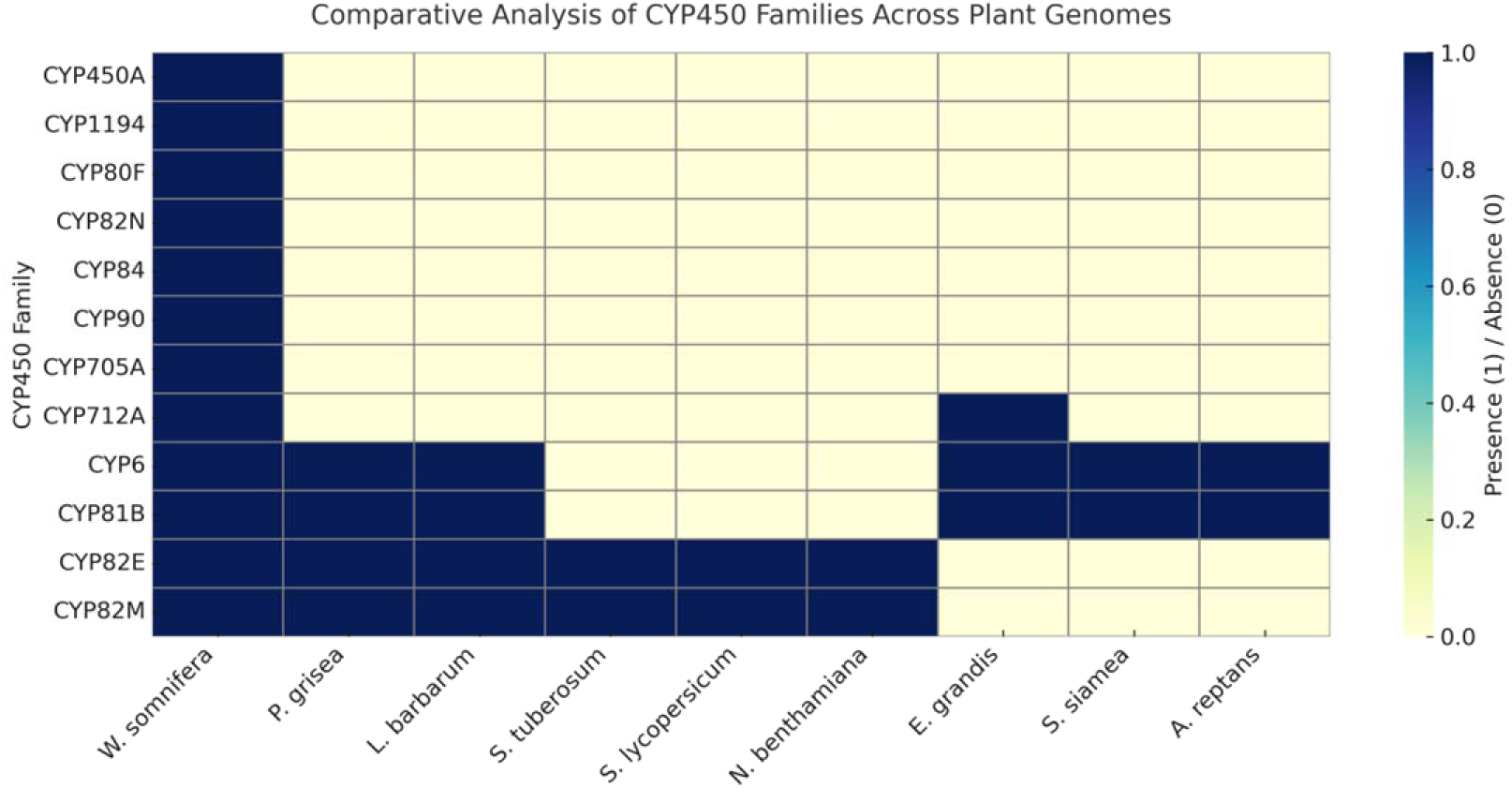
Comparative distribution of CYP450 families across selected plant genomes

**Table 3:** CYP450 gene families present in all nine selected plant genomes.

| Subfamily | *A.reptans* | *E.grandis* | *L.barbarum* | *N.benthamiana* | *P.grisera* | *S.tuberosum* | *S.Siamea* | *S.lycopersicum* | *W.somnifera* |
| --- | --- | --- | --- | --- | --- | --- | --- | --- | --- |
| **CYP51G** | 1 | 3 | 2 | 2 | 1 | 1 | 1 | 2 | 4 |
| **CYP701A** | 4 | 3 | 3 | 2 | 1 | 5 | 1 | 1 | 2 |
| **CYP704A** | 7 | 10 | 8 | 4 | 19 | 16 | 5 | 7 | 6 |
| **CYP707A** | 5 | 4 | 4 | 13 | 5 | 8 | 6 | 9 | 6 |
| **CYP710A** | 5 | 2 | 1 | 2 | 2 | 1 | 1 | 1 | 3 |
| **CYP714A** | 1 | 3 | 4 | 8 | 3 | 3 | 3 | 5 | 4 |
| **CYP716A** | 4 | 12 | 17 | 15 | 32 | 13 | 7 | 8 | 12 |
| **CYP718A** | 1 | 2 | 1 | 4 | 1 | 1 | 4 | 1 | 1 |
| **CYP71A** | 55 | 43 | 30 | 20 | 18 | 46 | 30 | 21 | 18 |
| **CYP71B** | 17 | 15 | 45 | 32 | 69 | 50 | 16 | 26 | 20 |
| **CYP721A** | 1 | 4 | 3 | 3 | 2 | 2 | 2 | 2 | 4 |
| **CYP72A** | 48 | 23 | 36 | 30 | 53 | 47 | 11 | 33 | 52 |
| **CYP734A** | 1 | 1 | 6 | 8 | 16 | 8 | 3 | 4 | 4 |
| **CYP735A** | 1 | 6 | 2 | 2 | 1 | 1 | 1 | 2 | 1 |
| **CYP73A** | 7 | 2 | 3 | 7 | 4 | 2 | 3 | 3 | 7 |
| **CYP74A** | 1 | 1 | 10 | 10 | 6 | 4 | 5 | 6 | 4 |
| **CYP75B** | 4 | 4 | 17 | 17 | 28 | 18 | 10 | 16 | 9 |
| **CYP76G** | 1 | 2 | 8 | 8 | 25 | 9 | 4 | 6 | 8 |
| **CYP77A** | 2 | 1 | 4 | 5 | 3 | 2 | 1 | 2 | 10 |
| **CYP77B** | 1 | 1 | 1 | 2 | 2 | 1 | 1 | 1 | 1 |
| **CYP78A** | 7 | 10 | 7 | 12 | 7 | 7 | 9 | 6 | 7 |
| **CYP79A** | 4 | 16 | 2 | 3 | 7 | 4 | 5 | 3 | 4 |
| **CYP82C** | 47 | 15 | 17 | 13 | 23 | 17 | 8 | 10 | 19 |
| **CYP84A** | 3 | 2 | 2 | 5 | 12 | 7 | 5 | 6 | 7 |
| **CYP86A** | 9 | 4 | 5 | 6 | 3 | 4 | 4 | 3 | 3 |
| **CYP86B** | 2 | 1 | 1 | 2 | 2 | 1 | 1 | 1 | 3 |
| **CYP88A** | 5 | 11 | 12 | 5 | 12 | 10 | 10 | 6 | 5 |
| **CYP89A** | 1 | 13 | 4 | 10 | 9 | 6 | 8 | 4 | 7 |
| **CYP90A** | 1 | 3 | 1 | 2 | 2 | 1 | 2 | 1 | 2 |
| **CYP94B** | 2 | 7 | 8 | 14 | 7 | 8 | 6 | 7 | 8 |
| **CYP94C** | 3 | 4 | 4 | 8 | 5 | 4 | 4 | 4 | 6 |
| **CYP94D** | 3 | 3 | 2 | 4 | 2 | 2 | 4 | 2 | 1 |
| **CYP97A** | 1 | 1 | 1 | 2 | 1 | 2 | 1 | 1 | 4 |
| **CYP97B** | 1 | 1 | 1 | 2 | 1 | 2 | 1 | 1 | 3 |
| **CYP97C** | 1 | 1 | 1 | 3 | 1 | 1 | 1 | 1 | 6 |
| **CYP98A** | 3 | 5 | 12 | 8 | 8 | 14 | 1 | 5 | 6 |

### 3.5 Gene Family expansion and contraction patterns by CAFE Analysis

Genome-wide CAFE analysis was carried out to highlight extensive expansion and contraction patterns across all nine species (Fig. 7). The results revealed that *N. benthamiana* exhibited the largest gene family expansion (+7841) alongside considerable contraction (−2223), reflecting its highly duplicated and dynamic genome structure. *W. somnifera* also showed substantial expansion (+4179) but an even larger contraction (−4889), suggesting an ongoing process of gene birth-and-death shaping its genome. Similarly, *L. barbarum* showed extensive expansion (+4688) with moderate contraction (−2347). Within the Solanaceae, *S. lycopersicum* (+2802 / -1564) and *S. tuberosum* (+2008 / -2140) both exhibited marked patterns of gene acquisition and loss, while *P. grisea* (+1412 / -4683) showed a strong bias toward contraction, indicating genome streamlining. Among non-Solanaceae, *E. grandis* (+3960 / -4593) and *S. siamea* (+2223 / -5414) both displayed extensive gene loss relative to gains, indicating a divergent course of genome evolution compared to *N. benthamiana* and *L. barbarum*.

**Fig. 7.**
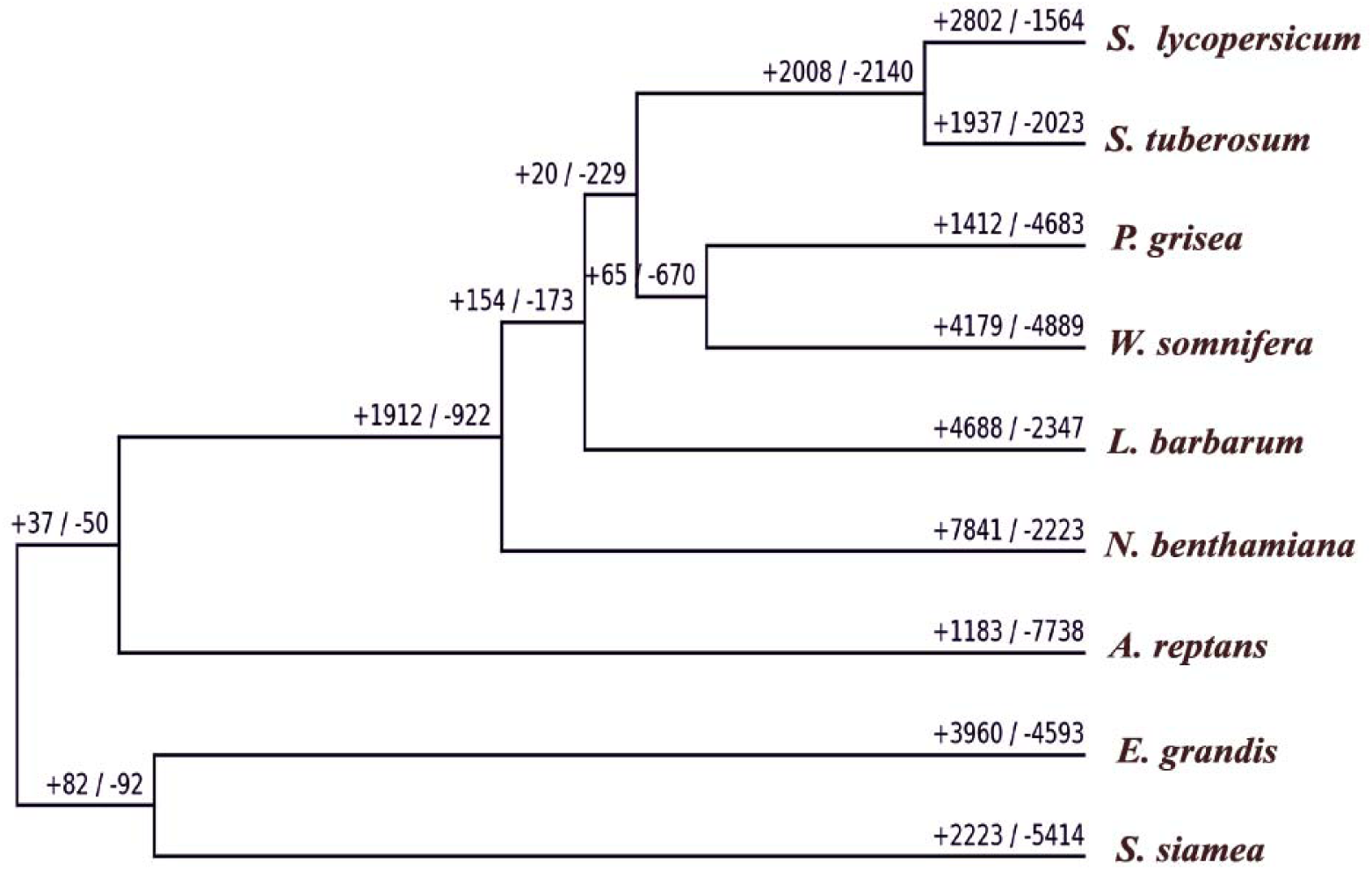
Phylogenetic tree illustrating all the gene families expansion (+) and contraction (–) counts across nine plant genomes

Since cytochrome P450 (CYP) gene families are well known for their tendency to undergo duplication and diversification, a focused CAFE analysis of the CYPome was carried out across the same nine species (Fig. 8). The CYPome analysis revealed distinct but interconnected evolutionary patterns. The strongest expansions were observed in *E. grandis* (+61) followed by *N. benthamiana* (+51), and *P. grisea* (+36), reflecting extensive diversification of CYP450 genes. Moderate expansion was observed in *S. tuberosum* (+30) and *W. somnifera* (+20), while *S. lycopersicum* showed only a small gain (+9). Contraction trends followed a different order, with *W. somnifera* showing the highest loss (–51), followed by *S. lycopersicum* (–28), *N. benthamiana* (–23), and *P. grisea* (–22). By contrast, *E. grandis* lost relatively few genes (–17) despite its large expansion.

**Fig. 8.**
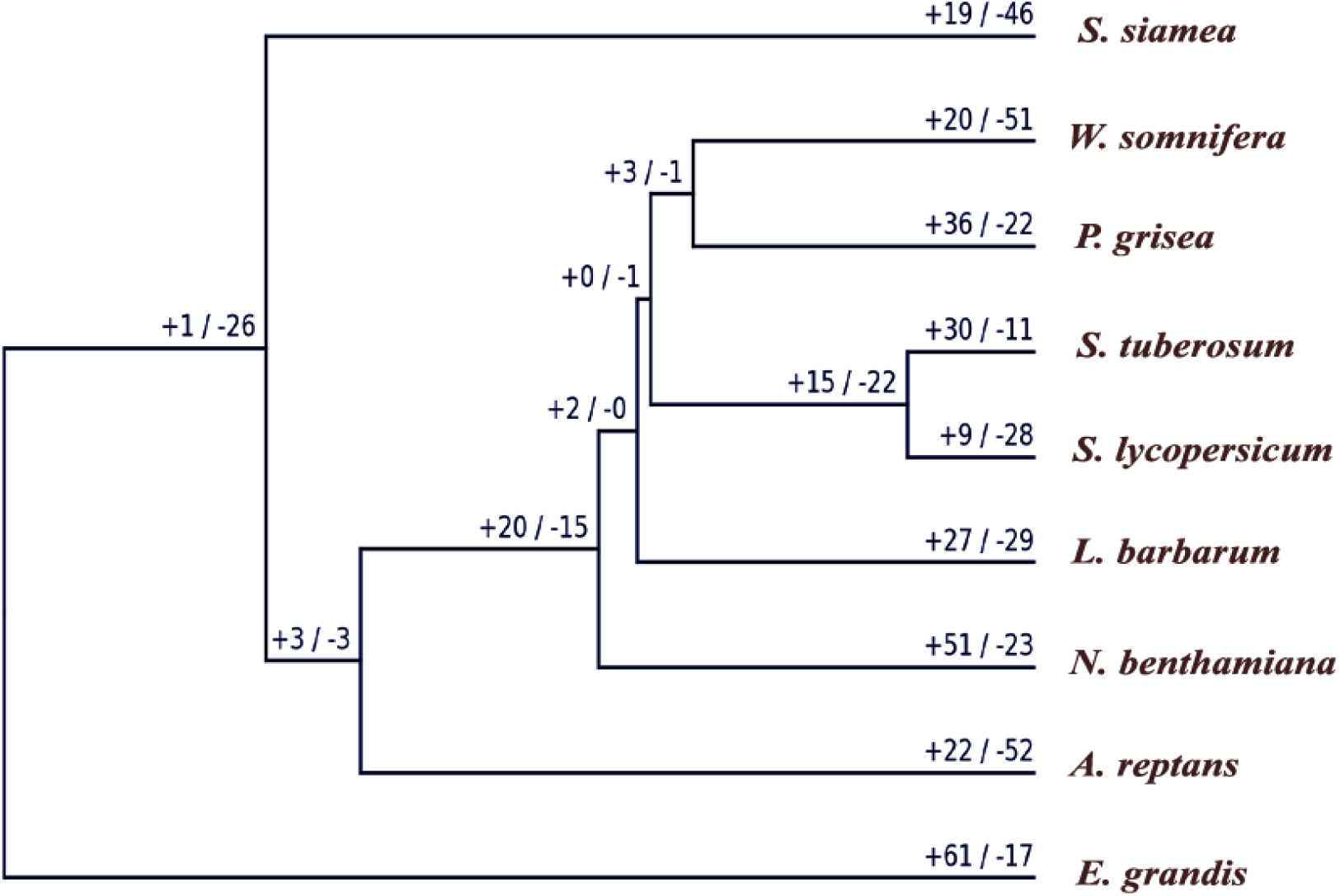
Phylogenetic tree illustrating all the CYP450 gene family expansion (+) and contraction (–) counts across nine plant genomes

Functional annotation of the CAFE enrichment analysis revealed that out of 113 identified CYP450 families, only 17 showed significant expansion or contraction patterns across the plant genomes studied. In *W. somnifera*, four CAFE families (0G000003, 0G000004, 0G000007, and 0G000010) exhibited significant contraction (Supplementary file 3). These families correspond to CYP subfamilies including CYP83B1, CYP76B1, CYP736A247, and CYP76A. Among them, CYP83B1 is known for its role in glucosinolate biosynthesis and auxin regulation in plants like *Arabidopsis*. CYP76B1 and CYP736A247 participates in detoxification of xenobiotics and cyanogenic glucoside biosynthesis, while CYP76A enzymes catalyze critical transformations in the biosynthesis of various terpenoids, including monoterpenoid oxidation a key step in producing monoterpenes involved in plant defense and signaling.

In addition to the notable CYP450 family contractions observed in *W. somnifera*, several other selected plant genomes displayed significant expansions or contractions in specific CYP subfamilies. For instance, *P. grisea* exhibited extensive expansions in families such as CYP71D55, CYP83B1, multiple CYP72A subfamilies (e.g., CYP72A554, CYP72A556, CYP72A692), and CYP736A, reflecting a broad amplification of metabolic capacity. Similarly, *S. tuberosum* showed expansions in CYP83B1, CYP76B6, CYP72A556, CYP71A4, and CYP88C1, while *A. reptans* and *E. grandis* demonstrated considerable gains in CYP81E, CYP82D, CYP76B, CYP81E22, and CYP749A families. Conversely, species such as *Solanum lycopersicum* and *Nicotiana benthamiana* presented selective patterns of expansion and contraction (Table 4).

**Table 4:** CAFE enrichment analysis.

| Genome | CYP Subfamilies with Expansion (+) / Contraction (-) |
| --- | --- |
| *Physalis grisea* | CYP71D55 (+40), CYP83B1 (+24), CYP81D11 (+17), CYP72A (+19), CYP736A (+17), CYP76A (+24), CYP82A (+9), CYP704C1 (+16) |
| *Solanum tuberosum* | CYP83B1 (+21), CYP76B6 (+21), CYP72A (+15), CYP71A4 (+18), CYP704C1 (+10), CYP76C2 (+9), CYP88C1 (+8), CYP75B1 (+13) |
| *Withania somnifera* | CYP83B1 (-6), CYP76B1 (-2), CYP736A (-3), CYP76A (-3) |
| *Nicotiana benthamiana* | CYP83B1 (-5), CYP72A (-4), CYP71A4 (-2), CYP72A (+8) |
| *Solanum lycopersicum* | CYP72A (-7), CYP93A3 (+) |
| *Ajuga reptans* | CYP81E (+28), CYP72A (+16), CYP82D (+47), CYP76B10 (+35) |
| *Eucalyptus grandis* | CYP81E (+28), CYP749A (+25), CYP82A (+20) |
| *Lycium barbarum* | CYP71A4 (+14) |

## Discussion

*W. somnifera* is renowned for its medicinal withanolides, yet limited genomic resources and the incomplete characterization of their biosynthetic pathway have constrained both functional insights and the use of genome editing for pathway improvement(Ahmad et al., 2024; Reynolds et al., 2024).Contextually, a high-quality genome assembly would provide a foundational resourceto facilitate pathway discovery, and establish a basis for future genetic improvement and metabolic engineering strategies. In the present study, the genome assembly of *W. somnifera* was generated using a hybrid approach combining Oxford Nanopore Technologies (ONT) long reads and Illumina short reads, resulting in a scaffold-level assembly estimated at 2.2 Gb with solid coverage and gene annotation completeness. Such a combined sequencing strategy involves the long-read capability of ONT to resolve complex genomic regions and structural variation, while Illumina reads improve base-level accuracy and error correction(Espinosa et al., 2024; Wang et al., 2025). Recent studies have demonstrated that this hybrid methodology enhances genome assembly contiguity and accuracy for complex plant genomes, enabling more reliable downstream functional genomics and breeding applications(Dumschott et al., 2020; Usha et al., 2022). The present assembly, while more fragmented than the reference reported by(Hakim et al., 2025) still captures the majority of coding regions, as reflected in its BUSCO completeness. The superior contiguity and completeness of the reference are largely due to higher long-read coverage and optimized assembly strategies. Nonetheless, our assembly provides valuable complementary understandings, particularly through the integration of short- and long-read data that enhance accuracy and coverage of repetitive regions. Comparative analysis between the two assemblies further revealed moderate levels of SNP and indel variation, indicating intraspecific genetic diversity that may contribute to phenotypic differences, especially in genes linked to secondary metabolite biosynthesis. Despite the availability of high-quality genome assemblies for *W. somnifera*(Hakim et al., 2025), the present study is significant as it highlights cultivar-specific genomic features. Chemotype-level assemblies are particularly valuable as they provide details into sequence polymorphisms and structural variations that contribute to metabolic diversity and local adaptation(Hämälä et al., 2021). This has been shown in other plant systems, including *Arabidopsis thaliana*, where reference genomes and hundreds of sequenced accessions exist, yet additional cultivar assemblies continue to exhibit genotype-specific variations with functional consequences(Lian et al., 2024). Similarly, studies in *Solanum* have demonstrated that assemblies from individual accessions can identify unique gene content, presence/absence variants, and structural rearrangements important for agronomic performance and metabolite production (Barchi et al., 2021; Molitor et al., 2024). Moreover, identifying unique sequence polymorphisms enhances the precision of genome editing. In CRISPR-Cas9 applications, SNPs in gRNA target sites can alter editing efficiency and specificity. Relying on a single reference genome may therefore increase the risk of off-target effects(Saha et al., 2025; Wang et al., 2020). Therefore, this genome assembly of a local *W. somnifera*(AGB-002) variety provides a strong foundation for functional genomics, metabolic engineering, and the molecular breeding of medicinally important traits.

Gene prediction from the *in-house* genome identified numerous non-redundant genes, with functional annotation showing roles in both primary metabolism and secondary metabolite biosynthesis. KEGG analysis revealed strong enrichment in isoprenoid and other secondary metabolite pathways, which are essential for terpenoid and steroid production and support the plant’s ability to produce bioactive compounds such as withanolides(Senthil et al., 2015). While primary metabolic pathways are fundamental across plant species due to their essential cellular functions, the clear presence of isoprenoid and secondary metabolite pathways among the top categories highlights the metabolic versatility of *W. somnifera*.

The present study further reports the identification of CYP450s from the genome of *W. somnifera*, which catalyze oxidative modifications critical for withanolide biosynthesis by shaping structural diversity and bioactivity(Shilpashree et al., 2022). While numerous studies have characterized CYP450s in other Solanaceae members(Hao et al., 2022; Lan et al., 2025), this represents the first comprehensive catalog for *W. somnifera* Phylogenetic analysis using experimentally validated triterpenoid-related CYP450s as references enabled the identification of potential candidates(Ghosh, 2017; Yasumoto et al., 2017). Herein, *W. somnifera* CYP450s showed clear subfamily-level clustering, validating their annotation and revealing several candidates closely associated with CYPs known to act on triterpenoid scaffolds.Specifically, *W. somnifera* CYP72A members (CYP72A692_1 and CYP72A560_4) clustered with *M. truncatula* CYP72A63 and CYP72A67, which catalyze C-30 and C-2 oxidations of β-amyrin (Biazzi et al., 2015; Fanani et al., 2019). Likewise, CYP716A48 grouped with *M. truncatula* CYP716A12, responsible for C-28 oxidation in oleanolic acid biosynthesis(Carelli et al., 2011), while CYP724B2 aligned with *A. thaliana* CYP708A2, linked to scaffold hydroxylation and steroid metabolism(Ghosh, 2017). In addition, CYP51G1 clustered with A. *strigosa* CYP51H10, an enzyme catalyzing β-amyrin epoxidation and hydroxylation in avenacin biosynthesis (Kunii et al., 2012). These associations indicate that *W. somnifera* has likely adapted orthologs or paralogs from conserved CYP subfamilies to catalyze analogous oxidative transformations including oxidation, hydroxylation, and epoxidation on triterpenoid or steroid scaffolds. By tailoring these backbones, such enzymes could play central roles in withanolide biosynthesis. Therefore, CYP72A692_1, CYP72A560_4, CYP716A48, CYP724B2, and CYP51G1 emerge as strong candidates for functional validation and pathway elucidation, offering critical entry points for understanding and engineering withanolide metabolism.

Since withanolides are present in certain Solanaceae species, absent in others, and even reported in a few non-Solanaceae plants it can be presumed that variation in CYP450s, may contribute to this scattered distribution(Dhar et al., 2015; Knoch et al., 2018; Narayanan and Nagegowda, 2024). Therefore, the present study entails a comparative analysis of the CYPome across nine selected plant genomes to explore whether differences in CYP450 gene content could influence withanolide biosynthetic potential. Although specific marker CYP450s distinguishing withanolide-producing from non-producing genomes were not identified, the study revealed distinct CYP450 genes unique to the *W. somnifera* CYPome. For example, the presence of rarely reported families such as CYP450A and CYP1194, which are uncharacterized in the plant groups, highlights possible species-specific gene innovation or horizontal gene transfer, reflecting adaptive evolution in response to unique metabolic demands(Hansen et al., 2021). CYP80F and CYP82N families are involved in the biosynthesis and modification of secondary metabolites, including alkaloids, flavonoids, and phenolics(Yonekura-Sakakibara et al., 2019; Zhan et al., 2022)while CYP84 and CYP90 play key roles in phenylpropanoid and brassinosteroid biosynthesis, respectively (Bancosİ et al., 2002; Meyer et al., 1996). Their occurrence exclusively in the *W. somnifera* genome, as compared to the other analyzed species, might be the result of adaptive evolution in response to ecological and metabolic demands. Similarly, the distinct presence of CYP705A, associated with triterpenoid diversification(Ghosh, 2017), suggests that *W. somnifera* may have adapted to expand triterpenoid metabolism, contributing to withanolide diversification.Further, it was observed that CYP81B and CYP6 families were absent particularly from the Solanaceae species (such as tomato, potato, and tobacco) thereby signifying taxonomically distinct differences in cytochrome P450 repertoires(Vasav and Barvkar, 2019). Since CYP81B enzymes have been reported in *Arabidopsis* to participate in indole glucosinolate metabolism and contribute to flavonoid modification and defense responses (Bak et al., 2001) their absence from certain plant families might signify functional replacement by other CYP450 families or reduced reliance on these pathways. Similarly, the occurrence of CYP6 in plants is unusual, as this family is primarily linked to xenobiotic detoxification in insects and fungi(Guo et al., 2016; Peng et al., 2017). Its presence may reflect rare horizontal transfer events or limited, lineage-specific functions(Ono et al., 2025). Furthermore, it was found that CYP82E and CYP82M were present only in Solanaceae genomes and absent from non-Solanaceae species such as *E. grandis, S. siamea, and A. reptans*. This restricted occurrence suggests their recruitment for clade-specific alkaloid biosynthesis, particularly in nicotine and tropane pathways characteristic of Solanaceae. Their absence in other lineages indicates an independent evolution of alkaloid metabolism, consistent with the distinct chemical profiles among these groups(Shoji et al., 2024; Zhang et al., 2023).

In addition to the unique CYP450s identified in the selected genomes the analysis also revealed a conserved set of 36 CYP450 subfamilies shared across all nine genomes, reflecting the strong conservation of core CYP450 functions. Although these CYPs were present in varying copy numbers across species, such differences are likely shaped by ploidy, gene duplication, and lineage-specific expansion(Fang et al., 2024). Representative sequences from each genome and subfamily were used to construct a phylogenetic tree, which resolved into distinct clades consistent with established plant CYP450 phylogeny. Within these clades, most CYPs formed orthologous subclades, with 243 of 250 sequences retaining expected relationships, highlighting their stable biochemical roles. Families such as CYP82C, CYP84A, CYP94A, CYP734A, CYP716A, and CYP78A showed strong conservation across species, suggesting their preserved roles in core metabolic processes, including triterpenoid and flavonoid modifications(Ghosh, 2017).Beyond this conserved core, several CYP450 subfamilies such as CYP75B, CYP71A, CYP71B, CYP716A, and CYP734A were displaced from their expected positions, suggesting diversification through duplication and adaptive recruitment into new metabolic roles(Chakraborty et al., 2023; Fang et al., 2024). Among them *W. somnifera* CYP75B and CYP734A, known for their roles in flavonoid, steroidal, and triterpenoid metabolismin other plant species (Hamberger and Bak, 2013; Hausjell et al., 2022; Ohnishi et al., 2006) might have diversified through duplication and were recruited for withanolide biosynthesis. These examples illustrate how evolutionary flexibility in CYP450s allows neofunctionalization, providing the raw material for specialized metabolic pathways(Hamberger and Bak, 2013; Nelson and WerckLReichhart, 2011; Rieseberg et al., 2023). Additionally, several triterpenoid-related CYP450 subfamilies such as CYP71A, CYP72A, CYP93A, CYP76A, CYP94B, CYP81E, CYP88A, and CYP79D were found to be shared across all nine genomes. (Fanani et al., 2019; Ghosh, 2017; Helliwell et al., 2001; Li et al., 2022; Nelson and WerckLReichhart, 2011). These enzymes catalyze oxidative tailoring reactions such as hydroxylation, epoxidation, and side-chain modifications, enabling the diversification of triterpenoid scaffolds(Ghosh, 2017; Yang et al., 2024). Their universal presence across distantly related species indicates that plants have retained common enzymes for diverse triterpenoids even though the final metabolites differ. This further suggests that structurally similar metabolites can be produced through diverse biosynthetic routes, where different CYP450s converge on comparable scaffold modifications(Limones-Mendez et al., 2020; Ono and Murata, 2023). In this regard, the occurrence of shared triterpenoid CYPs in both withanolide-producing Solanaceae and non-Solanaceae genomes indicate that while the complete withanolide pathway is lineage-specific, the core oxidative framework is widely conserved and adaptable for specialized metabolism.

Furthermore, CAFE analysis was conducted to investigate genome-wide gene family evolution, with a focus on cytochrome P450 (CYP450) families across nine species. The results revealed that CYPome evolution aligns closely with genome-wide patterns across the selected genomes. These variations are expectedly driven by genomic events such as segmental, tandem, and whole-genome duplications (WGDs) as is evident from the previous reports. For example, tandem duplication is a major driver of CYP450 family expansion in *Citrus clementina*, contributing about 41% of duplicated genes(Liu et al., 2023). WGDs also promote CYP450 expansion and functional diversification in terpenoid biosynthesis, as reported in *Solanum lycopersicum* (Tang et al., 2024), *Lavandula angustifolia*(Li et al., 2021), and *Ipomoea batatas* (Xing et al., 2022). In addition, tandem and segmental duplications have significantly contributed to the expansion of the CYP71 clan, a key lineage involved in secondary metabolism, as observed in *Capsicum annuum*(Hao et al., 2022). Considering these examples, the observed variation in CYPome and overall gene family sizes in *N. Benthamiana* (expansion) and *W. somnifera and P. grisea* (contractions) may be attributed to such duplication events that likely enhance chemical diversity and adaptability to ecological pressures. Conversely, species with relatively stable CYPomes, such as *S. tuberosum* and *S. siamea*, maintain a conserved CYP450 repertoire. This suggests that their existing gene complement sufficiently supports essential metabolic and ecological functions, thereby reducing selective pressure for extensive family expansion or contraction. (Fang et al., 2024).Further CAFE enrichment analysis revealed distinct patterns of CYP subfamily expansion and contraction across species. In *Withania somnifera*, which exhibited an overall reduction in its CYPome, several subfamilies such as CYP83B1, CYP76B1, CYP736A247, and CYP76A showed marked contraction. Since CYP76A is linked to the monoterpenoid pathway, this reduction might reflect a reallocation of metabolic resources favouring triterpenoid (withanolide) biosynthesis in this genome. These observations align with broader trends in plant metabolic evolution, where gene family contraction often accompanies pathway specialization and adaptive genome streamlining, leading to more efficient and specialized metabolic functions (Hansen et al., 2021; Kariñho-Betancourt et al., 2022).

Conclusively, this study presents the first high-quality genome assembly of the Indian *Withania somnifera* cultivar AGB-002, providing a platform for future research in functional genomics and genome editing. It offers a detailed profiling of CYP450 candidates in *W. somnifera* and other selected plant genomes, thereby enhances our understanding of how CYP450s contribute to metabolic evolution in the species. While a few promising candidate genes have been identified for their potential roles in triterpenoid biosynthesis, their functions still need to be confirmed through experimental studies. A chromosome-level assembly, which is not yet available, would make it possible to locate CYP genes precisely and explore how they are organized within triterpenoid biosynthetic clusters. Such mapping would help identify groups of co-localized CYPs and speed up efforts to decipher the withanolide pathway. Consequently, these results provide a framework for targeted gene validation, precise genome editing, and metabolic studies aimed at enhancing withanolide production and supporting the sustainable use of this medicinal plant.

## Acknowledgements

SG acknowledges the Department of Biotechnology, Government of India, for the fellowship as DBT-RA-II (DBT-RA/2023-24/N/IIIMJ/03). The authors gratefully acknowledge the Director of CSIR–Indian Institute of Integrative Medicine for providing research facilities and infrastructure. We also thank Negenome Biosolutions Pvt. Ltd. for their valuable support.

## Data availability

The sequences generated using hybrid platform of Illumina sequence data and Oxford Nanopore technology (ONT) have been submitted to NCBI sequence read archive (SRA) with accession number PRJNA1345563.

## Authors Contributions

S. Gupta conceived and designed the study, carried out data acquisition and analysis, interpreted the results, prepared all figures and tables, drafted the initial manuscript, and critically reviewed and approved the final version. P.Misra contributed to the interpretation of results and critically reviewed and refined the manuscript. R. Singh contributed to study conception and assisted in preparing figures and supporting materials. MK. Dhar guided the study design and interpretation and critically reviewed and approved the final manuscript. All authors read and approved the final manuscript.

## Code availability

Not applicable.

## Declarations

### Ethics approval

Not applicable.

### Consent to participate

Not applicable.

### Consent for publication

Not applicable.

## Figure and File legends

**Supplementary Fig. 1 Phylogenetic tree constructed using representative CYP450 protein sequences from each genome and subfamily.**

**Supplementary table 1: Shared Triterpenoid associated CYP450 gene families present in all nine selected plant genomes**

**Supplementary file 1 (.xlsx) List of cytochrome P450 (CYP450) genes identified from the nine selected plant genomes. The file includes gene identifiers, family and subfamily classifications.**

**Supplementary file 2 (.xlsx) Matrix showing the distribution of CYP450 families, subfamilies, and clans across the nine selected plant genomes. The file also highlights CYPs commonly shared among all genomes, including triterpenoid-related CYPs conserved across species.**

**Supplemetary file 3 (.xlsx) CAFE enrichment analysis highlighting 17 CYP450 families showing significant expansion or contraction across the nine plant genomes. Significantly affected families are marked in yellow**

