## Supplementary table 1 for "Whole-Genome sequencing of Indigenous *Withania somnifera* accession and comparative cytochrome P450 phylogenomics"

**Supplementary table 1: Shared Triterpenoid associated CYP450 gene families present in all nine selected plant genomes**

| CYP450 Subfamily | Clan | W. somnifera | A. rapens | E. grandis | Barbatu | Bentham | P. grisea | Senna | S. lycopersicum | S. tuberosum |
| --- | --- | --- | --- | --- | --- | --- | --- | --- | --- | --- |
| CYP71A | CYP71 | 18 | 43 | 30 | 20 | 18 | 46 | 30 | 21 | 18 |
| CYP72A | CYP72 | 52 | 23 | 36 | 30 | 53 | 47 | 11 | 33 | 52 |
| CYP93A | CYP71 | 6 | 8 | 5 | 7 | 10 | 4 | 25 | 4 | 6 |
| CYP76A | CYP71 | 8 | 31 | 11 | 15 | 28 | 4 | 9 | 16 | 11 |
| CYP94B | CYP86 | 2 | 2 | 6 | 7 | 13 | 6 | 2 | 1 | 3 |
| CYP81E | CYP71 | 36 | 9 | 7 | 12 | 12 | 8 | 5 | 5 | 5 |
| CYP88A | CYP85 | 5 | 11 | 12 | 5 | 12 | 10 | 10 | 6 | 5 |
| CYP79D | CYP71 | 1 | 1 | 1 | 2 | 1 | 1 | 1 | 1 | 2 |
