## Supplementary figures and images for "Whole-Genome sequencing of Indigenous *Withania somnifera* accession and comparative cytochrome P450 phylogenomics"

### Supplementary Fig 1

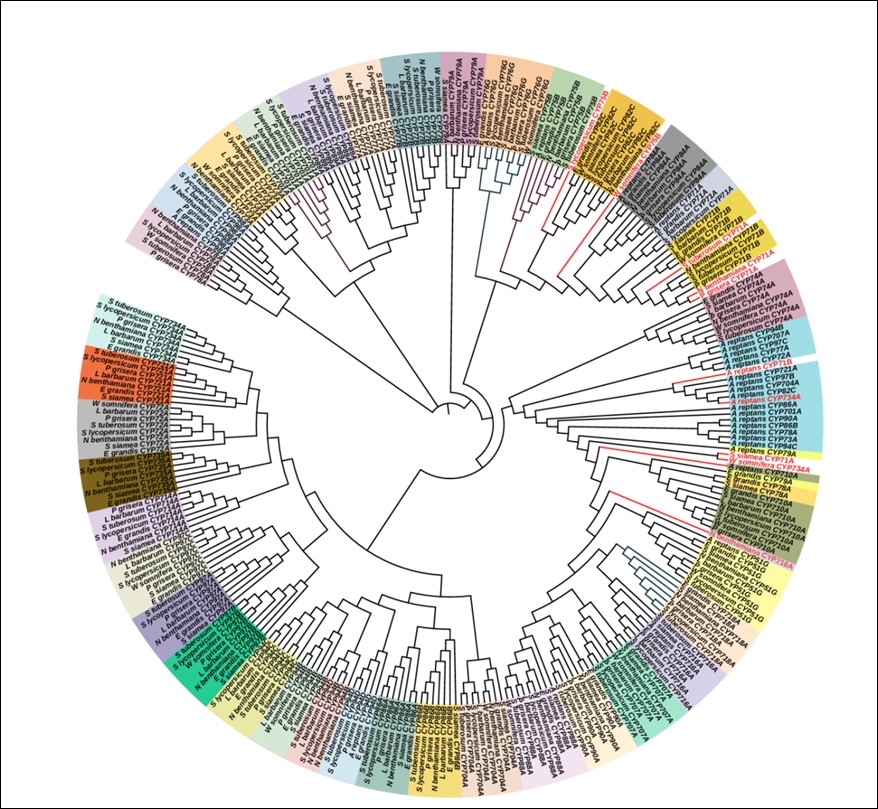
